# Social Status as a Latent Variable in the Amygdala of Observers of Social Interactions

**DOI:** 10.1101/2024.07.15.603487

**Authors:** SeungHyun Lee, Ueli Rutishauser, Katalin M. Gothard

**Affiliations:** Department of Neurosurgery, Massachusetts General Hospital, Harvard Medical School, Boston, MA, USA; Department of Neurosurgery, Cedars-Sinai Medical Center, Los Angeles, CA, USA; Computation and Neural Systems, Division of Biology and Biological Engineering, California Institute of Technology, Pasadena, CA, USA; Department of Physiology, College of Medicine, The University of Arizona, Tucson, AZ, USA

**Keywords:** Macaques, facial expressions, dPCA, scanpath, fixations, context, mixed selectivity

## Abstract

Successful integration into a hierarchical social group requires knowledge of the status of each individual and of the rules that govern social interactions within the group. In species that lack morphological indicators of status, social status can be inferred by observing the signals exchanged between individuals. We simulated social interactions between macaques by juxtaposing videos of aggressive and appeasing displays where two individuals appeared in each other’s line of sight and their displays were timed to suggest the reciprocation of dominant and subordinate signals. Viewers of these videos successfully inferred the social status of the interacting characters. Dominant individuals attracted more social attention from viewers even when they were not engaged in social displays. Neurons in the viewers’ amygdala signaled the status of both the attended (fixated) and the unattended individuals suggesting that in third party observers of social interactions, the amygdala signals jointly the status of interacting parties.

**Highlights:**

- Monkeys infer the social status of conspecifics from videos of simulated dyadic interactions
- During fixations neural populations signal the social status of the attended individuals
- Neurons in the amygdala jointly encode the status of interacting individuals

**In brief:** Third party-viewers of pairwise dominant-subordinate interactions infer social status from the observed behaviors. Neurons in the amygdala are tuned to the inferred dominant/subordinate status of both individuals.

## Introduction

High social status is one of the most valued and coveted social commodities by both humans and non-human primates as it ensures access to resources, reproductive success, social support, and ultimately better health and well-being.^1, 2, 3, 4, 5, 6^ The desire to rise in a hierarchy or retain one’s status directly or indirectly shapes all interactions within a social group.^7, 8, 9, 10, 11, 12^ These interactions rest on a mutual understanding of social rules and norms that govern relationships. By observing hierarchical interactions, social primates learn to accurately place each member of their social group into a specific position on the social ladder, forming a linear hierarchy.^13, 14, 15, 16, 17^

Macaque societies are distributed on a continuum between egalitarian, tolerant societies (e.g., Tonkean macaques) where status plays a minor role in social interactions, and despotic societies where strict linear hierarchies are strongly enforced.^18, 19^ Rhesus monkeys live in despotic societies^20, 21^ where status is established based on aggression and coalition-building skills in males, and the rank of the mother and birth order in females.^22, 23^ Dominance is expressed through aggressive (agonistic) behaviors toward subordinates, whereas low status is expressed through submissive, appeasing displays.^13, 24, 25^ Thus, this species is ideally suited to explore the social-cognitive and neural foundations of social organization.

We investigated whether rhesus macaques can infer a social hierarchy from observing videos of simulated interactions among unfamiliar individuals. Given that high status is not exteriorized by morphological markers such as skin or fur coloration in rhesus monkeys,^26^ the viewer monkeys were expected to infer status from observing pairwise interactions among “video monkeys” organized by design into an arbitrary linear hierarchy. To determine whether the behaviors in the videos elicited neural responses encoding social status, we recorded and analyzed the activity of neurons in the amygdala of two viewer monkeys. We hypothesized and confirmed that the inferred social status of the observed individual manifests in status-related changes in neural activity.

## Results

### 1. Looking Time Proportional to the Status of the Observed Individuals

Two adult male monkeys, M_A_ and M_D_, watched 740 and 928 videos, respectively, depicting dominant-subordinate dyadic interactions among unfamiliar monkeys. During the trial, we simultaneously recorded the viewers’ scanpaths and neural activity (**Figure 1A**). **Figures 1B and 1C** illustrate that each hierarchy group consisted of four individuals, resulting in six pairwise interactions per group. We created four distinct hierarchy groups, totaling 24 videos. Each video was repeatedly watched, ranging 58-74 repetitions. To simulate natural interactions, we juxtaposed pairs of videos, designating one monkey as dominant and the other as subordinate. The two monkeys appeared to be in each other’s line of sight, and the timing of the threatening-appeasing displays was adjusted to simulate the natural cycle of dominant-subordinate signals exchanges.

**Figure 1.**
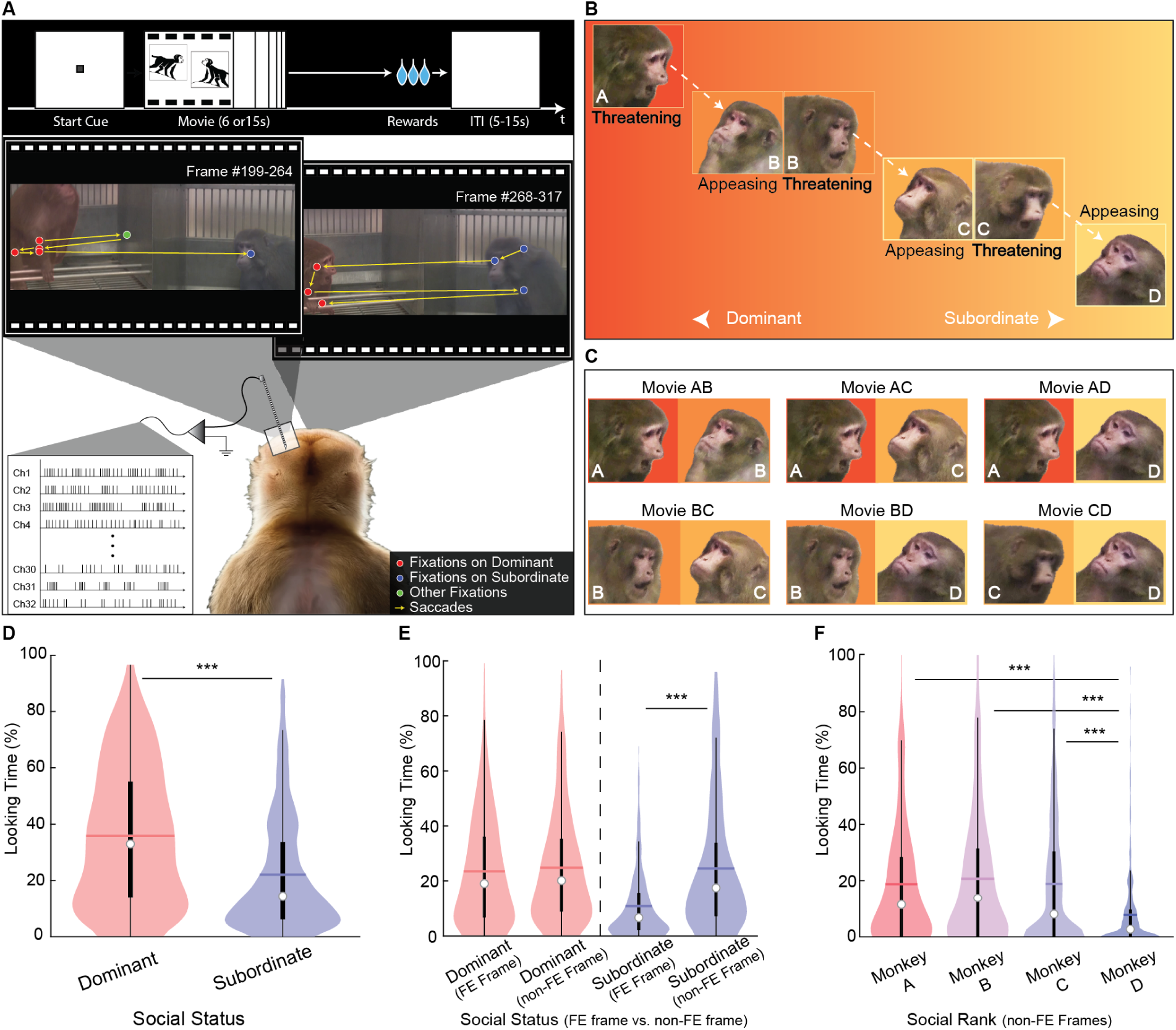
**(A)** Trial flow and experimental setup. Viewer monkeys with electrode arrays in their amygdala watched videos depicting simulated pairwise social interactions. Each trial began with the viewer fixating on a start cue and receiving reward for watching the videos. Eye tracker recorded scanpaths (sequences of fixations and saccades). In two example scanpaths segment, red and blue circles denote fixations on dominant and subordinate monkeys, respectively, while yellow arrows represent the saccades connecting these fixations. **(B)** An example linear hierarchy constructed from juxtaposed videos of aggressive and appeasing displays of four male monkeys (A, B, C, and D). **(C)** Pairwise interactions among monkeys A, B, C, and D. Monkey A consistently displayed threatening towards all partners, while monkey D consistently exhibited appeasing towards all partners. Monkeys B and C alternated between dominant and subordinate roles depending on their social partner. (**D**) Proportion of cumulative fixation duration on dominant and subordinate individuals relative to the total playing time of the movie (15s). (**E)** Proportion of looking time at frames depicting facial expressions (FE) and neutral faces (non-FE) relative to the total screen time of FE and non-FE frames. **(F)** Proportion of cumulative looking time at neutral faces of each hierarchical individual relative to the total screen time of frames with neutral faces.

We analyzed the average time viewers spent looking at each monkey in the dyad and compared the proportion of time spent looking at the dominant and subordinate animals in each trial (see **Star+Method** for calculation details in **Figures 1D-F**). On average, the viewer monkeys spent more time looking at the dominant animal (Mean: *μ* =35.87Hz, Standard Deviation: *∂* =24.18Hz) than the subordinate animal (*μ* =22.07Hz, *∂∂* =20.19Hz) in the dyad (**Figure 1D**; t-test, t(4534)=20.86, p=2.41×10^-92^). Notably, the threatening facial expressions produced by the dominant animals do not account for the longer looking times, as comparable looking times were elicited by threatening (*μ*=21.29Hz, *∂*=19.73Hz) or neutral facial expressions ( *μ* =21.00Hz, *∂* =21.56) of the dominant animal (red violin plots in **Figure 1E**; t-test, t(1501)=0.27, p=0.79). Moreover, viewers allocated less attention to subordinate animals when they displayed appeasing facial expressions (*μ*=9.01Hz, *∂*=11.53Hz) compared to neutral facial expressions (*μ*=14.43Hz, *∂*=23.46Hz) (blue violin plots in **Figure 1E**; t-test, t(1353)=-5.56, p=3.30×10^-8^), suggesting that viewers allocate attention to and gain information from both neutral and dominant/subordiante displays. Indeed, when comparing viewers’ looking times at the neutral faces of the four monkeys in the hierarchy, the least fixated neutral faces were those of the lowest-ranking monkey D (**Figure 1F**), while the differences among the other monkeys were not significant (one-way ANOVA with factor identity, df=3, F=21.38, ***: p<0.001, A vs. B: p=0.63, A vs. C: p=1.00, A vs. D: p=1.64×10^-9^, B vs. C: p=0.70, B vs. D: p=7.11×10^-13^, C vs. D: p=1.74×10^-9^). These findings collectively suggest that increased social attention (measured by total looking time at the dominant animal in a dyad) primarily reflects social status rather than mere their identity or facial expressions alone.

### 2. Neurons in the Amygdala Respond to Social Status of the Attended Individuals

We recorded neural activity of 202 well-isolated single neurons in the amygdala of two viewer monkeys. Our goal was to investigate whether the neural activity elicited by watching the videos carries information about the social status of the protagonists. Specifically, we compared the firing rates during fixations on the faces or bodies of dominant and subordinate monkeys. Prior to analyzing the contribution of social status on neural activity in the amygdala, we verified that these videos elicited neuronal responses that are comparable to previously documented responses to static images or videos of single individuals^27, 28^ and to multiple faces presented simultaneously.^29^ As expected, a two-way ANOVA analysis (factors: identity and facial expression, see **STAR+Methods**) replicated previous findings: 28.22% of amygdala neurons were selective for individual faces/bodies, fewer (7.92%) were selective for facial expressions, and a subclass of neurons (5.94%) showed mixed selectivity for both identity and facial expression.

**Figure 2** shows two example status-responsive neurons, recorded from the central (**Figures 2A-D**) and accessory basal (**Figures 2E-H**) nuclei of the amygdala. We analyzed firing rates during fixations on dominant versus subordinate individuals, independent of the fixation target’s identity and regardless of whether the previous fixation was on a dominant or subordinate. As the status of the mid-ranking monkeys B and C, depended on the social partner, we analyzed fixations separately according to each individual’s social status in each video pair. In the first example neuron, firing rates during fixations on dominant animals (*μμ*=11.00Hz, *∂∂*=9.19Hz) were significantly higher across all dominant animals in the six pairwise interactions (**Figure 2B**; t-test, t(856)=4.13, p=3.97×10^-5^) compared to fixations on subordinate animals (*μμ*=8.66 Hz, *∂∂*=7.39Hz). Conversely, in the second example neuron, firing rates during fixations on the subordinate animals (*μμ*=15.37Hz, *∂∂*=12.20Hz) were significantly higher (**Figure 2F**; t-test, t(655)=-3.16, p=1.70×10^-3^) than during fixations on dominant animals (*μμ*=12.49Hz, *∂∂*=11.05Hz). This was the case despite the great heterogeneity of fixation targets (various individuals, facial expressions, and face or body areas). The temporal patterns of status-related responses were analyzed using fixation-aligned histograms of firing rates, indicating time bins where the firing rates were significantly higher when viewing dominant animals (**Figure 2D**; using t-test with sliding window, bin size = 60 ms, stride size = 10 ms, *: p<0.05). Another histogram (**Figure 2H**; employing the same methods as in **Figure 2D**) highlighted time bins with significantly lower firing rates when fixating on dominant animals. Notably, fixations in **Figures 2C and 2G** can be preceded by fixations on either dominant or subordinate individuals, thus it was possible to detect significantly different firing rates in the time bins preceding the fixation onset (time 0). Nevertheless, both neurons exhibited tuning to the social status of the currently fixated individual. At the population level, 24.75% of recorded amygdala neurons (50 out of 202) responded with significant changes in firing rates to either dominant or subordinate status (**Figure 2I**). Neurons included in these analyses were recorded from all the major amygdala nuclei, predominantly localized in the basal (BA) and central (CeA) nuclei (**Figure 2J and K**).

**Figure 2.**
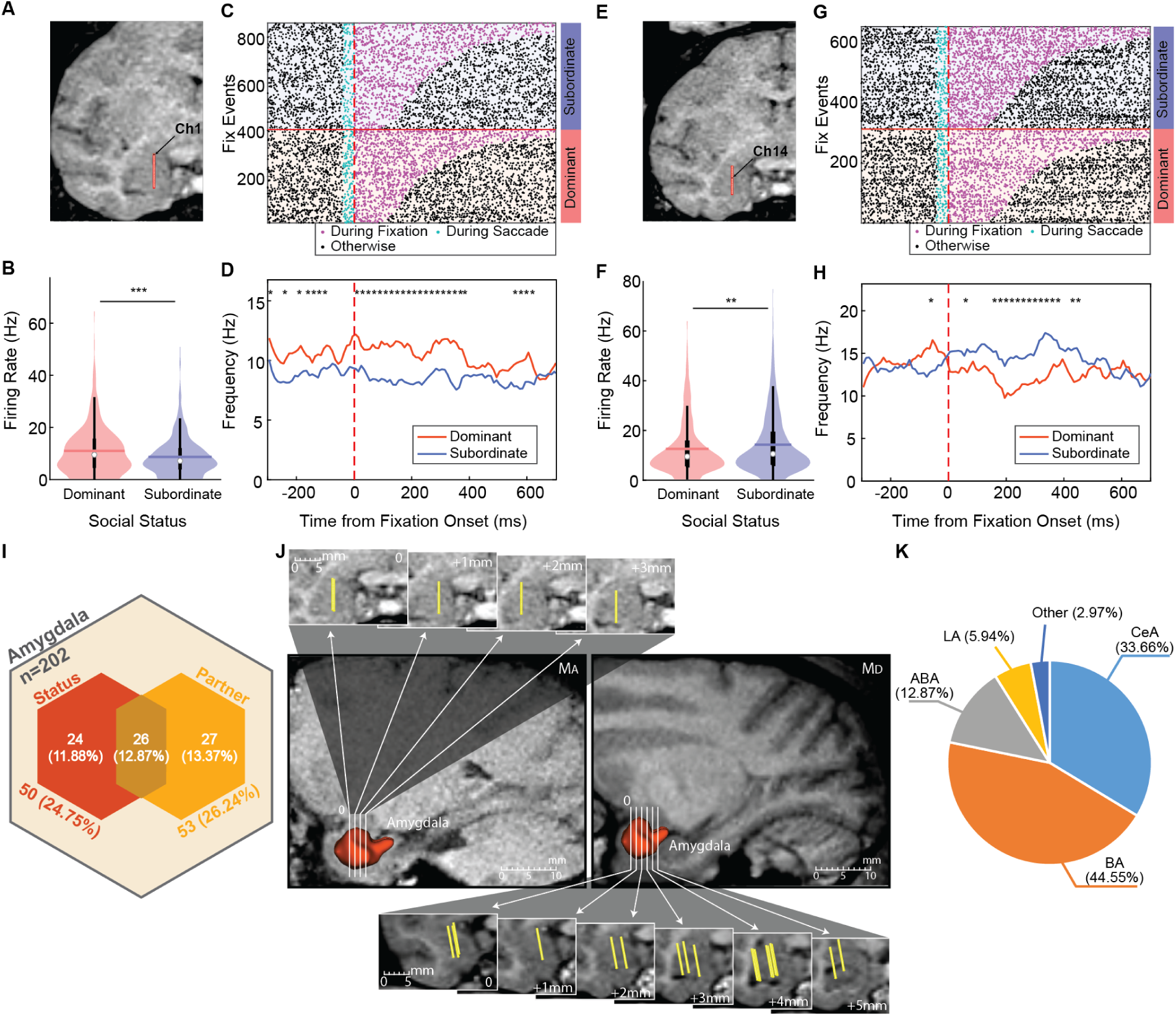
Example neurons tuned to the social status of fixated individuals. Coronal MRI slices in panels **A** and **E** indicate the position of a 32-contact V-probe in the amygdala, marked by red lines. Panels **A-D** depict the neuron recorded from channel 1 in the central nucleus, whereas panels **E-H** show an example neuron recorded from the channel 14 located in the basal nucleus. **(B)** Firing rates were significantly higher during fixations on dominant animals. **(C)** Raster plot showing fixations on subordinate (top) and dominant (bottom) animals in a dyad, sorted by fixation duration. Magenta dots represents spikes during fixations, and cyan dots represent spikes during preceding saccades (with saccade durations ranging 36-54 ms). **(D)** Temporal pattern of status-related responses in a fixation-aligned histogram of firing rates, indicating time bins with significantly higher firing rates during fixations on dominant animals (0-400 ms). **(F)** Firing rates during fixations on the subordinate animals were significantly higher. **(G)** Separate rasters for fixations on subordinate (top) and dominant (bottom) animals in a dyad, sorted by fixation duration. **(H)** Temporal pattern of status-related responses in a fixation-aligned histogram of firing rates, indicating time bins with significantly lower firing rates during fixations on dominant animals (0-400 ms). **(I)** Venn diagram showing the proportion of amygdala neurons tuned to social status and social context (partner effect). **(J)** Recording locations in M_A_ and M_D_, with yellow bars indicating electrode positions across all 23 recorded sites from 20 recording days. **(K)** Distribution ratio of collected amygdala neurons among different subregions (n=202).

Status emerges as a latent variable in neural activity, alongside manifest variables like face identity, indicating mixed selectivity. For monkeys, A and D, ranked highest and lowest in the hierarchy, identity and social status were not dissociable because their consistent dominance or subordination across all three partners. However, mid-ranking monkeys, B and C, could dissociate face identity from social status, as they alternated between dominant and subordinate roles depending on the partner (**Figures 1B and C**). A two-way ANOVA analysis (factors: social status and identity) of fixation-related firing rates while viewing monkeys B and C showed not only a main effect of status (16.83% of recorded neurons) but also a main effect of identity (20.79% of recorded neurons), along with a significant status-identity interaction (13.86% of recorded neurons). Given that close to 25% of recorded neurons showed tuning to social status, the small effects at the level of individual neurons may amount to large signals at the population level (**Figure 2I**). To test this hypothesis, we conducted a group analysis of all neurons.

### 3. Status-related population responses in the amygdala

We used demixed principal component analysis (dPCA)^30, 31^ to determine status-related changes in population firing rates during fixations on each monkey. This method, a supervised approach, captures the variance in population activity explained by two factors and their interaction: *Movie* (depicting the six dyads), and *Status* (dominant and subordinate). Across all components, *Status* (13%) and *Movie* (44%) explained the largest proportions of variance in neural activity, followed by their interaction (37%) (**Figure 3A**). This finding aligns with the observation that status-related responses varied with the social partner (partner effect), as each movie depicted a different pairing of four animals in the hierarchy. To incorporate fixations with varying durations into the dPCA model, we normalized each fixation duration to a unit time of 1 second, segmented into 20 time points. Indeed, dPCA component #1 (*Status*, red box, **Figure 3B**), contributes 6.9% of the explained variance and clearly separates normalized firing rates in response to looking at dominant (solid lines) versus subordinate (dotted lines) monkeys across all six movies.

**Figure 3.**
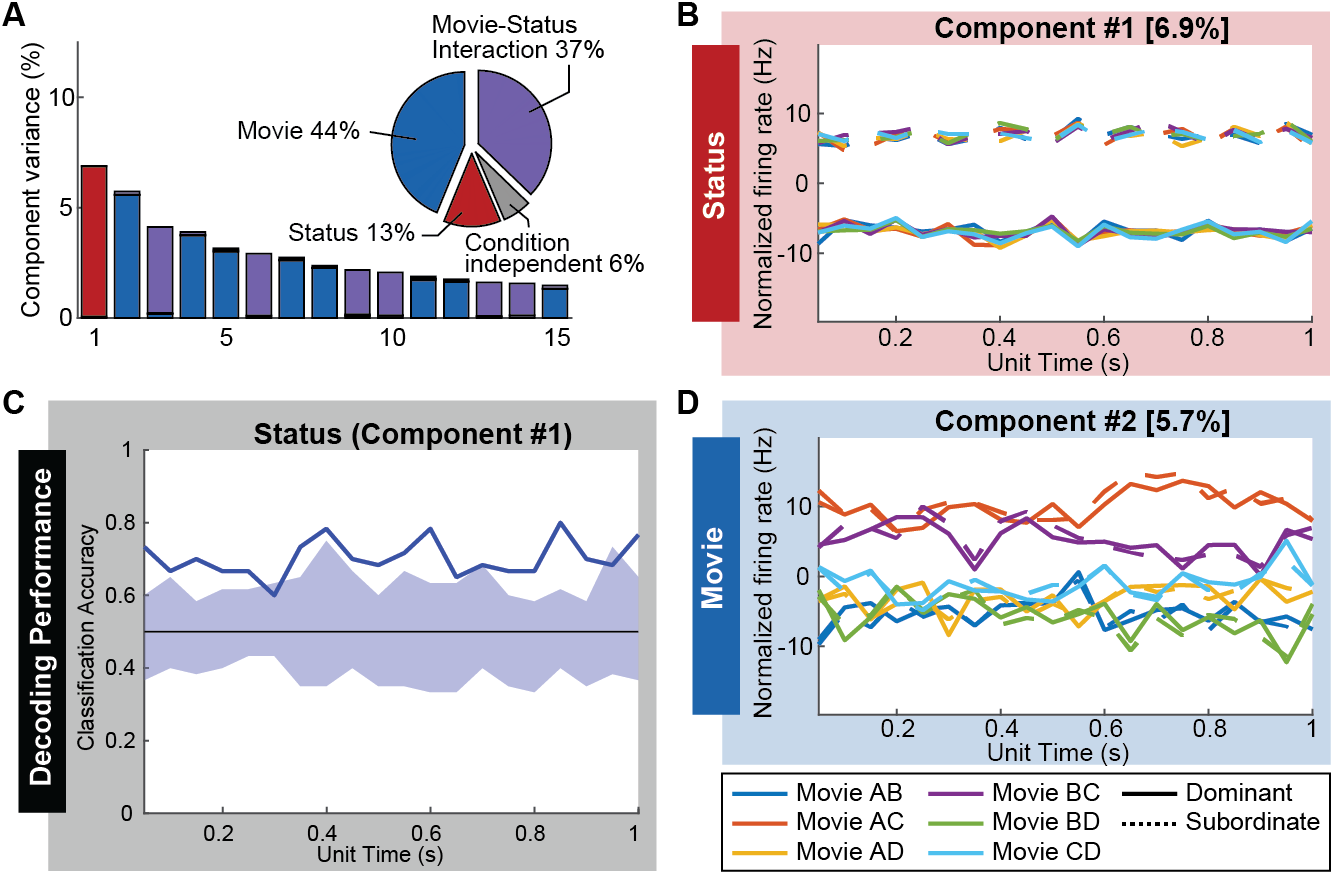
dPCA reveals the primacy of social status in neural population activity during fixations. **(A)** Ratio of variance contributed by each component and factor. Each “Movie” refers to one of the six distinct video clips featuring different dyads. **(B)** Normalized firing rate of the primary component (#1) representing “Social Status” during a unit time of the fixation, and its contribution to explained variance. **(C)** Classification accuracy of component #1, indicating that amygdala firing rates during fixations are influenced by the social status of the target, surpassing random chance. The shaded area represents the range of classification accuracy expected by chance, estimated from 100 shuffling iterations, while the solid line indicates the classification accuracy of component #1. **(D)** Normalized firing rates of higher-order components representing “Movie” type (#2) during a unit time of fixation.

Social status was decodable from the activity of dPCA component #1, as indicated by the classification accuracy (solid line in **Figure 3C**), which exceeded chance levels. This range was estimated from 100 iterations of a shuffling procedure, confirming the reliable presence of this signal in the amygdala. Component #2 (*Movie*, blue box, **Figure 3D**) contributes 5.7% of the explained variance and separates movies AC and BC (orange and purple lines) from the other four movies. In these composite movies, one side shows monkey C appeasing moneys A and B. Component #2 likely segregates the appeasing behavior specifically, rather than just the identity, of monkey C, as movie CD, which depicts monkey C threatening, does not separate out in this component. In contrast to social status (component #1), movie identity was not decodable at classification accuracy levels higher than chance (additional details in **Figure S1)**.

The dPCA results showed that beyond face identity and facial expression, the population of neurons in the amygdala, whether individually tuned to status or not, contains information about the status of interacting individuals and the specific video being watched. As a control, we repeated the dPCA on amygdala recordings while subjects watched movies of moving objects, formatted identically to the hierarchy movies. In this control condition, classification accuracy for “status” did not surpass chance levels when status was arbitrarily assigned to moving objects (**Figure S2**).

### 4. The partner effect: neural responses to the status of the attended individual are modulated by the status of the unattended individual

We asked whether the status-related response during fixation on a dominant or subordinate monkey was different when the partner was different. If the firing rates while looking at the same animal displaying the same behaviors are fully explained by the content of the viewed video, the identity of the social partner should not significantly alter the observed neural responses. Recall that the same videos were used to depict dominant or subordinate behaviors of each monkey in the hierarchy (the same video depicted monkey A as dominant to monkeys B, C and D). Thus, fixations on monkey A (the highest-ranking animal) are expected to elicit the same firing rates regardless of whether the partners are monkeys B, C, or D. Contrary this prediction, however, we found a “partner effect” in 26.2% of the 202 amygdala neurons (i.e., a significant main effect of social partner on the firing rate of the fixated individual, computed using one-way ANOVA). **Figure 4** shows two example neurons that illustrate this partner effect.

**Figure 4.**
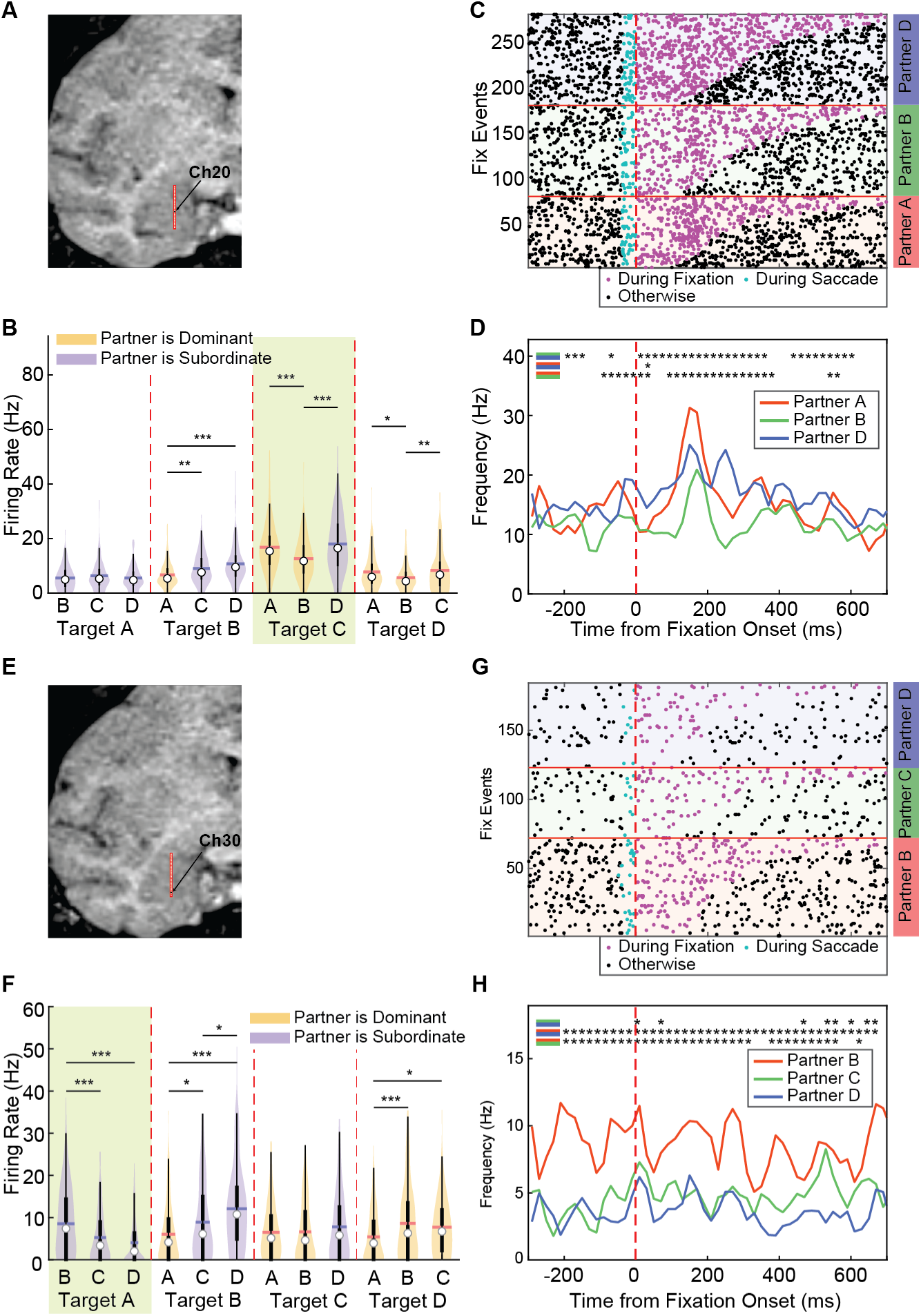
Neurons showing the partner effect. The red line superimposed on the coronal MRI slices in panels **A** and **E** indicates the position of a 32-channel V-probe in the amygdala and the closest contact to the example neuron. Panels **A-D** depict the neuron recorded from channel 20 in the accessary basal nucleus, whereas panels **E-H** show an example neuron recorded from the channel 30 located in the basal nucleus. **(B)** Each triplet of violin plots depicts firing rates during fixation on a monkey from the hierarchy (Target A, B, C, and D) with each possible partner. The first triplet represents firing rates during fixations on monkey A with partners B, C, and D. The second triplet (separated by red dotted lines) represents firing rates during fixations on monkey B with partners A, C, and D. Partner status relative to the fixated individual is color-coded: yellow for dominant and blue for subordinate. Bars and stars correspond to the statistically significant differences (One-way ANOVA, *p<0.05, **p<0.01, ***p<0.001). **(C)** Rasters of spike trains aligned to the onset of fixations on the middle-ranking monkey C (triplet with lime green background in panel **B**), paired with monkeys A (red), B (green), and D (blue). **(D)**. Fixation-aligned firing-rate histogram corresponding to the raster in panel **C**; using t-test between pairs of sliding windows as indicated by color legend in the upper left corner (bin size = 60 ms, stride size = 10 ms, *: p<0.05). **E-H**. The same plots for a neuron located in the basal nucleus. The violin plots with the lime green background corresponds to the triplet of fixating monkey A while paired with B, C, and D, as indicated by the red, green, and blue boxes and lines in panel **G** and **H**.

The neuron shown in **Figure 4A-D** exhibits a strong partner effect when looking at monkey C, where the firing rate increases significantly when the partner is D compared to B, and also increases when the partner is A compared to B (one-way ANOVA, df=2, F=9.07, ***: p<0.001). Furthermore, this neuron shows a strong fixation-related increase in neural activity at a latency of approximately 100 ms after fixation onset on monkey C, but this activity is significantly suppressed when the partner is monkey B (**Figure 4D**). In the second example neuron (**Figure 4E-H**), firing rates during fixation on monkey A are higher when the partner is B than C or D, despite all three partners displaying similar appeasing expressions. Note that the firing rate in response to fixating on monkey A decreases as the status of the social partner decreases (one-way ANOVA, df=2, F=10.42, ***: p<0.001). Likewise, firing rates during fixations on monkey D are lower when the partner is A compared to B and C.

The partner effect was presented in 53 out of 202 monitored neurons (26.2%), a proportion comparable to 24.7% neurons that showed a status effect (**Figure 2I**). Comparable proportion of neurons showed only a status effect, only a partner effect, and mixed selectivity for both (11.8%, 12.8% and 13.3% respectively). Overall, the partner effect suggests a joint representation of the two monkeys, where one animal serves as the focal point of the viewer’s social attention while the other provides the “social context” in which the viewed individual is processed. This reflects the complex interplay between social status and social interactions encoded in amygdala neuronal activity.

## Discussion

### Videos of simulated interactions are sufficient to inform the status-dependent allocation of social attention in macaques

We report that the viewers of dyadic interactions extracted the status of the observed individuals and expressed their knowledge by allocating increased social attention toward dominant individuals. Indeed, in various social tasks, both humans and macaques allocate more looking time to high-status individuals.^11, 12, 32, 33, 34, 35, 36, 37, 38^ The looking preference did not emerge simply from more fixations on facial expressions, as the dominant individuals received similar levels of attention when they were shown with neutral disposition. Likewise, the looking preference is not informed by visible features of status, such as face coloration in dominant mandrills.^39^ In rhesus macaques, these visible features are either unrelated^26^ or only weakly related to social status.^40, 41^ Despite the artificial nature of the stimuli, it is remarkable that the viewers extracted the status of each monkey. This suggests that for rhesus monkeys, understanding social status is important, and they can make use even incomplete information to gain this knowledge.

### Social status as a latent variable in neural activity

In response to the static images of faces, neurons in the amygdala of both humans and monkeys respond to variables such as face identity,^27, 29, 42, 43, 44^ facial expressions,^27^ and even the social status of familiar individuals within an established hierarchy known to the viewer.^45^ Our results are novel because the individuals depicted in our videos were not personally familiar to our subjects. Here, social status emerged as a latent variable based on the observed dominant-subordinate interactions, independent of identity and facial expression (see **Figure 1**). In a similar human study, the BOLD signal increased in the amygdala as participants learned the relative status of individuals in a hierarchy.^46^ Because one of our subjects (M_D_) was familiar with these videos by the time neural recordings started, we were unable to track the experience-dependent emergence of the status-related neural responses. M_D_ had participated in a prior study exploring gaze following and shared attention saccades elicited by these videos that were also status-related.^12^ In these experiments, M_D_ was more likely to produce joint-attention saccades from the eyes and faces of the subordinate monkeys, indicating a prior understanding of the hierarchical position of each animal.

Higher social status was not always associated with higher firing rates. As shown in **Figure 2E-H**, some neurons responded to dominant status with a suppression of firing rates. Response suppression as a form of stimulus selectivity has been well-documented in the amygdala (e.g., [47] Mosher et al., 2010) and is related to features of circuit architecture that support highly complex behaviors through disinhibition (e.g., [48] Han et al., 2017).

### Neurons in the amygdala represent jointly the status of interacting individuals

Our findings add a novel dimension to the role of the amygdala in social perception. Not only do individual neurons respond to the social status of the attended individual, but their responses also depend on the unattended individual, who serves as a context signal. The joint representation of the status of multiple individuals supports the idea that neurons in the monkey amygdala show mixed selectivity or multidimensional responses.^49, 50, 51, 52, 53, 54^ Indeed, the amygdala has been emerged as a key node for processing social status since the earliest studies in non-human primates. Without an intact amygdala, monkeys fall in the social hierarchy^55, 56, 57^ and fail to display status-appropriate submissive and dominant behaviors during interactions with lower or higher-ranking conspecifics.^58, 59, 60^

In an even broader framework, the amygdala encodes both the status of others and the status of self. Self-status is reflected in gray matter density in the amygdala^61^ and in fMRI signals.^62, 63^ A similar effect has been shown in humans: lower perceived socioeconomic status causes greater activation of the amygdala.^64^ Self-status may be driven by subjective experiences, such as gains and losses during confrontations, which shape an individual’s expectations for future interactions and prepare the individual to extend these expectations to social partners. In species with strictly linear hierarchies, the status of others is among the defining features of an individual, akin to age, sex, and physical appearance. However, while status is abstract, age and sex may be gleaned from physical appearance. The data presented in this paper is an initial step toward understanding the neural substrate for the abstract and latent representation of social status in the primate amygdala.

## Acknowledgements

Funded by: P50MH100023

We thank Tess Marie Champ, Derek O’Neill, Natalia Magnusson, and Dr. Blake Tyler Seaton for help with animal training and data collection. Michael A. Cardenas and Archer I. Bowman edited the manuscript.

## Autor Contribution

Experimental design: SL, KMG; Data collection: SL; Data analysis and interpretation, SL, KMG, UR; manuscript preparation: SL, KMG, UR.

## STAR+METHODS

### RESOURCE AVAILABILITY

#### Lead Contact

Further information and requests for resources and reagents should be directed to and will be fulfilled by the lead contact, Katalin M. Gothard.

#### Materials Availability

This study did not generate new unique reagents.

#### Data and Code Availability

The data that support the findings of this study are available from the lead contact upon reasonable request. All original code is publicly available as of the date of publication. Any additional information required to reanalyze the data reported in this paper is available from the lead contact upon request.

### SUBJECTS DETAILS

We recorded single-unit spike activities in the amygdala, hippocampus, medial prefrontal cortex (mPFC), and other closely located areas from two adult male rhesus macaques (M_A_: 10 years old, 13.6 kg; M_D_: 9 years old, 11.8 kg). Meanwhile, we tracked the scanpath of these subjects as they watched visual stimuli (hierarchy movies). A shortcoming of the study was the limited size of our dataset due to an abdominal emergency (unrelated to the experimental procedures) developed by M_A_. This animal was euthanized after only 5 recording sessions.

## METHODS

### Video Stimuli

Each side of video clips had a resolution of 640 x 480 VGA, and these clips were combined to create the impression of 6 hierarchical interactions (AB, AC, AD, BC, BD, and CD) among a group of 4 monkeys. In this paper, we used four male hierarchy groups (P1, P3, P4 and P5. Where P stands for patriline). The juxtaposed movies were displayed on the screen at a resolution of 1920 x 1080 FHD and played for 6 or 15 seconds at a frame rate of 25 frames per second, resulting in each movie consisting of 125 or 375 frames. We manually segmented each monkey’s facial and body area in all frames and manually scored the frames containing facial expressions. The number of pixels in the facial area compared to the body area is reported in **Table 1**. Additionally, the number of frames with and without facial expressions is reported in **Table 2**. Note that P4 and P5 were designed later than P1 and P3, with the intention of increasing the number of frames showing the animals with facial expressions. Therefore, we used only the looking time results from when the viewers watched the P1 and P3 hierarchies for analysis, as shown in **Figures 1D-F**.

**Tabel 1.**
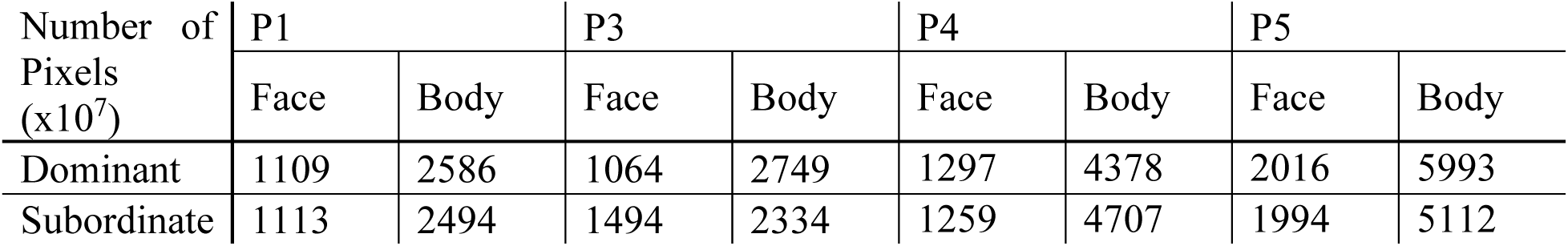
The number of pixels of the facial area and body area.

**Tabel 2.**
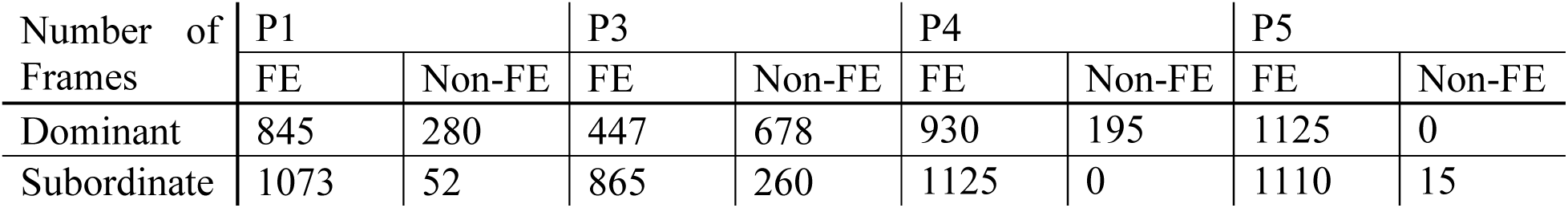
The number of frames with facial expressions and without facial expressions.

### Task flow

Video timing (stimulus onset and offset) triggered by the subject’s gaze on the starting cue was controlled using Monkey Logic (https://monkeylogic.nimh.nih.gov/). Each recording session (day) consists of several blocks, each containing 36 trials. Each trial began with a fixation of 250 ms on the 20 x 20 pixel-sized start cue, with a 100 x 100 pixel-sized invisible error boundary in the center of the monitor. There was no restriction for where the monkeys looked, the subject could freely scan the video or look away. After the video played (either 15 s or 6 s in duration), the subject received a small drop of juice reward (the same amount for all videos). The subject monkeys were not subjected to food or fluid restriction.

### Collecting Eye Tracking Data and Processing

Eye movements were recorded using an eye tracker (ETL-200, ISCAN Inc.) with a sampling rate of 120Hz. Eye tracker calibration was controlled by the hardware and software that provided by ISCAN Inc., and we calibrated the eye position at the beginning of each recording session. Eye tracking data was collected through voltage units, capturing both vertical and horizontal eye movements. To streamline analysis, we designated horizontal eye movement data as ‘Eye X’ and vertical data as ‘Eye Y’. As illustrated in **Figure S3**, raw data is initially recorded in millivolts (mV), then converted into pixel coordinates using calibration steps at the begging of every recording session. Our method for pinpointing saccade or fixation onset leverages the energy of the signal (*E*), as detailed in **equation (1)**, when *X* denotes the input signal, using n=30 as a window size. This proposed method highlights significant energy shifts, allowing us to distinguish between the saccadic and fixation periods. To determine the precise onset timing of saccades and fixations, we identify the onset by detecting when the second derivative of the energy signal crosses zero.

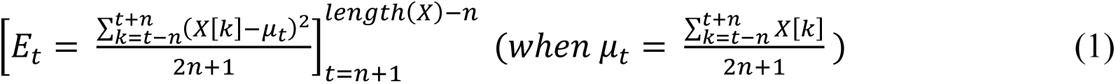

As mentioned, we used the eye tracking system with a sampling rate of 125Hz, indicating that the recording signal refreshes every 8 milliseconds on our data acquisition system (OmniPlex, Plexon Inc.), operates at a sampling rate of 1kHz. Since the eye tracking system has a lower resolution, there is a potential risk for timing inaccuracies of up to 8 milliseconds. To address this issue, we proposed the up-sampling approach outlined in **equation (2)**. This method proves highly effective in mitigating existing challenges. In our analysis, when *X* denotes the input signal, we implemented n=4 as a window size.

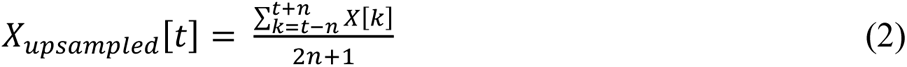

As depicted results in **Figure S4**, the distributions of both saccade duration and fixation duration exhibit greater uniformity, with reductions in artifacts attributed to the lower sample rate (up to 8ms timing error) evident across all recording days. To juxtapose the original and up-sampled results, we employed the Kolmogorov-Smirnov test (KS test, utilized the Matlab function kstest2). Days where the p-value from the test falls below 0.05 are denoted with an asterisk (*). Given that saccade durations typically fall within the range of 20-60 ms, shorter than fixation durations spanning 60-600 ms, the impact of up-sampling is notably more pronounced in saccade duration distributions. This is reflected in the KS test results, demonstrating significantly different for all recording days in the saccade duration plots (**Figure S4A**).

### Equations for the Looking Time Analysis

We measured the proportion of the looking time in **Figures 1D-F**. As below, we defined the equations for each panel to calculate the proper proportion of the looking time for each trial.

In **Figure 1D**, 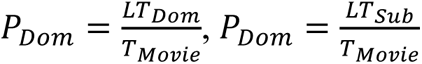

In **Figure 1E**, 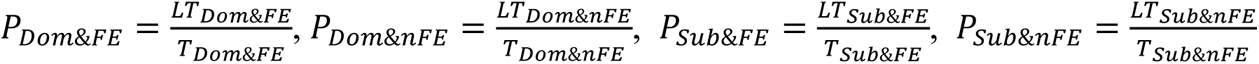

In **Figure 1F**, 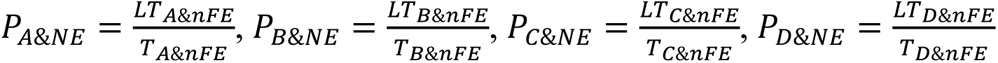

(When, *P* = *Proportion of looking time*, *LT* = *Looking Time, T* = *Playing Time*)

### Single-unit Spike Recording and Identification of Stable Cells

Single-unit spiking activity was recorded using a linear electrode array (V-probe, Plexon Inc.) with 32 channels spaced at 200 µm intervals along a 260 µm diameter shaft. The target recording locations were determined based on pre-scanned magnetic resonance imaging (MRI) slices of each subject. We used a motorized micro-drive system (Thomas Motorized Electrode Manipulator (MEM), Thomas Recording), which has a 1 µm axial resolution, to advance the electrode.

We utilized KiloSort for spike sorting, following the methodology outlined by the literature.^65^ For each sorted cell, we generated a cell specification sheet containing waveform data, inter-spike interval (ISI) histograms, spike count, ISI histogram stretched to the overall recording duration, and instantaneous firing rate throughout the recording session. Using a comprehensive list of criteria, we assessed whether each spike train was suitable for further data analysis. For example, we assessed the stability of the instantaneous firing rate over the recording duration and filter out neurons with an average firing rate below 0.5Hz during task sessions. Also, neurons with fluctuating firing rates over a block of movie trials were eliminated. If a cell specification meets all criteria, we designate the neuron as a “good cell” and include it in further data analysis. In total, we collected “good cells” from 202 neurons in the amygdala (M_A_: 106, M_D_: 96).

### Visualizing Data Set Distributions

To visualize the distribution and probability density of each data set, we employed violin plots, which are hybrids of a box plots and kernel density plots. This method offers the advantage of depicting both summary statistics and the density of each variable. In our violin plots, the inner thick black box represents the interquartile range, while the inner thin black line expanding this box denotes 1.5 times the interquartile range. The white dot signifies the median point of the variables. The shaded area illustrates the probability density of the data variables, with thicker part indicating higher probability.

### Quantifying Cell Responses Tuned to Social Status and Partner Effect

We classified cells that exhibited a significant difference in firing rates 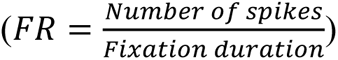when looking at dominant versus subordinate individuals as tuned to social status (see **Statistics** below). Similarly, if at least one of an attended target’s partners showed a significant difference in firing rates compared to the other two partners, we referred to this type of cell as tuned to the partner (partner effect). The PSTH plots in **Figures 2 and 4**, we used a 10 ms bin for the spike counts for the histogram. We smoothed the signals in the PSTH plots solely for the purposes of better visualization, using Savitzky-Golay filtering (Matlab function *sgolayfilt*) with a polynomial order of 3 and a frame length of 11. Note that, we calculated statistics before applying the smoothing to the signals.

### Statistics

In the PSTH plots, we used a t-test to assess the significant levels when comparing signals within the sliding window. The window size was 60 ms, and the stride size was 10 ms.

We used one-way and two-way ANOVA to statistically compare the effect of different on the firing rate of individual neurons. Significance levels (p-value) below 0.05, 0.01, and 0.001 are denoted with one, two, and three asterisks (*, **, ***), respectively.

We ran a one-way ANOVA to test for single factors. For example, we used it to compare looking times based on social status, the presence of facial expressions, facial area versus body area, and hierarchy rank. Additionally, we counted the number of neurons that were selective to the attended target’s social status or partner based on the results of the one-way ANOVA.

We utilized two-way ANOVA to explore how various factors affect neural activity in combination. To show the effects of the attended target’s identity and facial expressions, we conducted a two-way ANOVA with the factors of identity (monkey A, B, C, and D) and facial expressions (presence or absence of facial expressions). We also used a two-way ANOVA with the factors of social status (dominant and subordinate) and facial expressions (presence or absence of facial expressions) to test if the facial expressions of the fixated target affect neural activity instead of social status. Lastly, we tested for the main effect of status (dominant and subordinate), in addition to effect of identity (monkey B and C), using two-way ANOVA. For both one-way and two-way ANOVA, we ran the analyses independently for each neuron to test the effect on neural activity.

To compute dPCA results, we adapted Matlab package provided by literature^31^ (https://github.com/machenslab/dPCA) and customized its configurations for use with our datasets. We tested classification accuracy for 20 components, arranging them in descending order based on the explained variance for each marginalization. The classification accuracy value is derived from the 100 shuffling iterations. In the figures, we showed only the decoding accuracy plot corresponding to the highest-contributing component in the status factor results.

## Supplementary Figures

**Figure S1.**
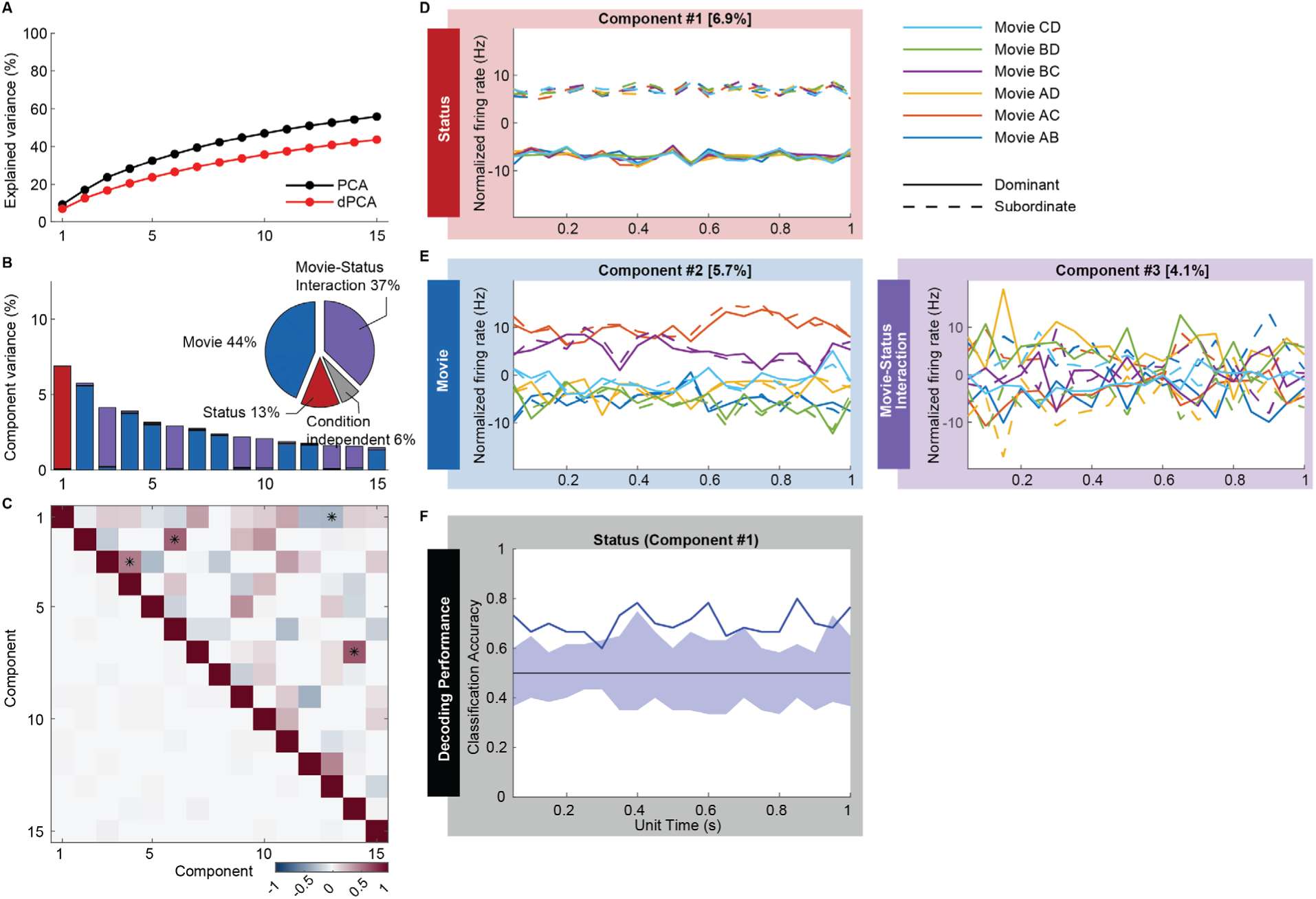
dPCA results of amygdala neurons during the hierarchy task. Detailed figure of Figure 5. (A) Cumulative (across the 15 components) explained variance comparing PCA and dPCA. (B) Ratio of contributed component variance of each component and factor. Movie refers to one of the six video clips each, with a different pairing of four animals in a hierarchy. (C) Dot products between demixed principal axes, with stars marking pairs in non-orthogonal (upper triangle). Correlations between demixed principal components (bottom triangle). (D) Normalized firing rate of the highest order component (#1) representing the factor of social status during a unit time of the fixation and its contribution to explained variance. (E) Normalized firing rates of the high-order components representing the factor of movie type (#2) and movie-status interaction (#3) during a unit time of the fixation. (F) Classification accuracy of component #1, indicating that firing rates during fixations in the amygdala are influenced by the social status of the fixation target, surpassing random chance.

**Figure S2.**
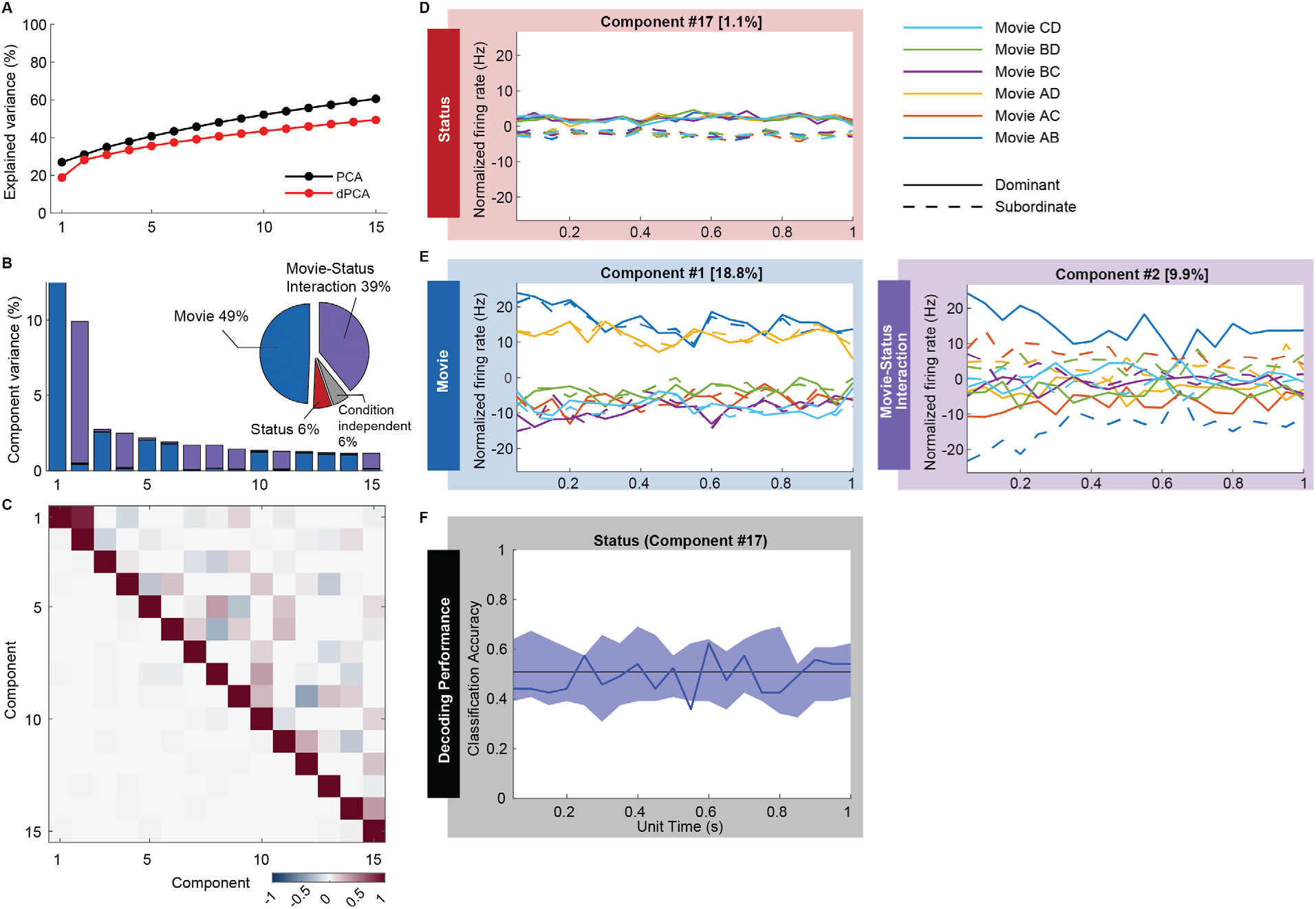
dPCA result of amygdala neurons during object task (control experiment). (A) Cumulative explained variance comparing PCA and dPCA. (B) Ratio of contributed component variance of each component and factor. Movie refers to one of the six video clips each, with a different pairing of four animals in a hierarchy. (C) *Dot products between demixed principal axes, with stars marking pairs in non-orthogonal (upper triangle). Correlations between demixed principal components (bottom triangle).* (D) Normalized firing rate of the component #17 representing the factor of social status during a unit time of the fixation and its contribution to explained variance. (E) Normalized firing rates of the components representing the factor of movie type (#1) and movie-status interaction (#2) during a unit time of the fixation. (F) Classification accuracy of component #17, indicating that firing rates during fixations in the amygdala are not influenced by the meaningless status of the fixation target.

**Figure S3.**
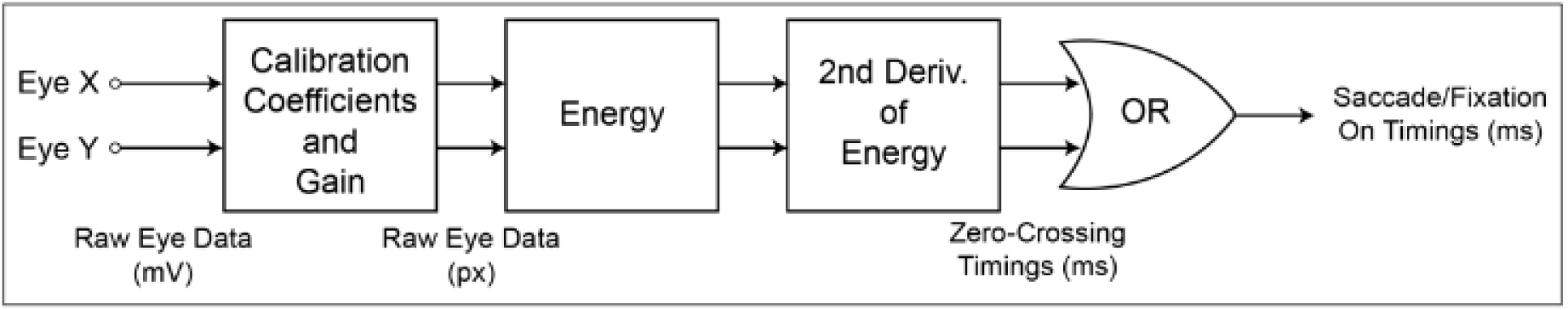
The flow chart of the process of the eye tracking data enhancement.

**Figure S4.**
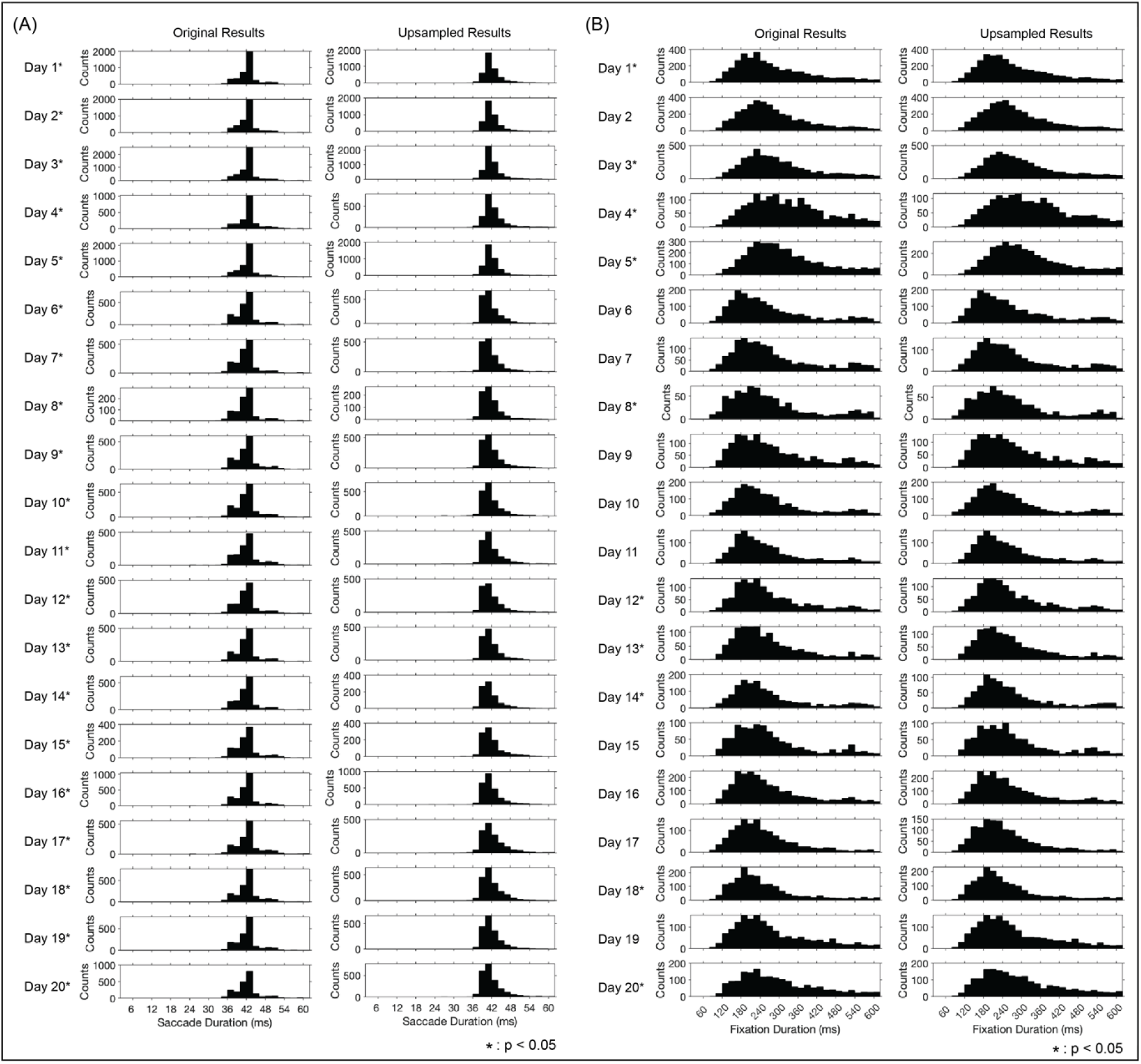
The results of improvements through the up-sampling. (A) Results in the distribution of the saccade duration. (B) Results in the distribution of the fixation duration.

